# Complex microbial consortia improve yield and physiological performance of leafy greens under deficit irrigation

**DOI:** 10.64898/2026.04.05.716566

**Authors:** Anna Edlund, Virginie Boreux, Abhinav Grama, Josh L. Espinoza, Shib Sankar Basu, Jamison McCorrison, Jack A. Gilbert, Thomas W. Crowther

## Abstract

Water scarcity is an increasing constraint on agricultural productivity, particularly for high-value vegetable crops. Deficit irrigation can reduce water use but frequently compromises yield and crop physiological status. We evaluated five microbial consortia (A–E) developed as products from a genome-sequenced culture collection. The consortia were dominated by spore-forming *Streptomyces* and Bacillales, four also contained mycorrhizal fungi, and their compositions reflected functional prioritization as well as manufacturing and regulatory constraints. Consortia were applied at two rates to soil-grown lettuce (*Lactuca sativa*) and spinach (*Spinacia oleracea*) in four greenhouse trials under full irrigation or a 30% deficit. Deficit irrigation reduced yield of untreated plants by approximately 10% (lettuce) and 13% (spinach). Relative to untreated deficit-irrigated controls, consortia increased yield by 2–8% in lettuce and 4–12% in spinach, significantly in five and six of ten treatments, respectively; most treatments still yielded less than fully irrigated controls. All treatments reduced wilting incidence and Water Band Index, and most reduced harvest delay and increased root length. Higher rates tended to produce stronger responses. Inoculant persistence and effects on resident soil communities were not measured, so mechanisms remain to be resolved.

**Sustainability Statement:** This work is primarily associated with SDG 6 (Clean Water and Sanitation), showing that complex microbial consortia can partially offset the yield losses of leafy greens grown under 30% deficit irrigation, contributing to agricultural water-use efficiency. By reducing the yield penalty associated with lower irrigation input, microbial inoculants offer a biologically derived tool for more sustainable water management in intensive vegetable production. This work is also relevant to SDG 2 (Zero Hunger), by supporting food production resilience under climate-driven water stress; SDG 13 (Climate Action), through adaptation strategies for water-limited agriculture; and SDG 15 (Life on Land), by reintroducing beneficial soil microorganisms into agricultural soils.

## Introduction

Agricultural productivity is increasingly constrained by water scarcity, soil degradation, and climate variability, with drought emerging as a primary limitation across global cropping systems (Daryanto et al., 2016; IPCC, 2023; Lesk et al., 2016). Reduced water availability disrupts essential plant physiological processes, including photosynthesis, nutrient uptake, and growth (Blum, 2010). Leafy vegetables represent roughly 18% of the EU’s fresh vegetable cultivation area according to the EU’s statistical overview reports in 2024 (Eurostat, 2026). This is a sector facing intensifying irrigation pressure as water scarcity deepens across European growing regions. In southern Europe, including Spain, a large share of lettuce and spinach is grown in soil, either in open fields or in soil-bound protected cultivation, although soilless production of leafy greens in controlled-environment systems is expanding. Soilless systems can recirculate irrigation water and support root microbiomes that differ from those in soil (Thomas et al., 2023), so both the relevance of deficit irrigation and the design of microbial inoculants depend on the production system. Identifying sustainable strategies for enhancing yield resilience of soil-grown crops under drying conditions is critical for offsetting future yield losses under climate change scenarios.

In addition to climatic drivers, soil structure is a primary regulator of plant-available water, as aggregation, porosity, and organic matter content regulate water retention and root access to moisture under stress (Six et al., 2002; Tisdall & Oades, 1982). Soil microorganisms play a central role in regulating the soil–plant water interface, influencing both soil physical properties and plant responses to water stress through a range of mechanisms (Ngumbi & Kloepper, 2016; Vurukonda et al., 2016). For example, microbial production of extracellular polymeric substances (EPS) promotes soil aggregation and can enhance pore connectivity and water retention (Chenu, 1993; Costa et al., 2018). Symbiotic fungi and filamentous soil bacteria also extend the effective root system through hyphal networks, increasing plant access to water and nutrients beyond rhizosphere depletion zones (Nadeem et al., 2014). Yet, intensive agricultural management has been associated with reduced soil biodiversity across European soils (Tsiafouli et al., 2015), which may diminish these services. As such, a growing body of research suggests that the reintroduction of beneficial microorganisms into degraded agricultural soils has the potential to enhance yield resilience.

Most studies exploring the potential of microbial inoculants to support plant physiology during stress have focused on single-or dual-species formulations, which may perform inconsistently across heterogeneous soil environments and variable stress conditions (Ocallaghan et al., 2022). In contrast, natural soil microbiomes function as complex, interacting communities (Zhalnina et al., 2018). This has motivated the development of defined microbial communities (a.k.a. SynComs) and other complex multi-species inoculants that more closely reflect the functional diversity of natural microbial systems (Alzate Zuluaga et al., 2024; Coker et al., 2022; Zhang, Gilbert, et al., 2025; Zhang, Jing, et al., 2025). Conceptual frameworks for defined microbial community design range from reductionist communities used to establish causality in plant–microbe interactions to trait-based communities designed for crop resilience (Vorholt et al., 2017; de Souza et al., 2020; Shayanthan et al., 2022). Designed communities have produced predictable plant phenotypes under nutrient limitation (Herrera Paredes et al., 2018), conferred salt-stress resilience to tomato in the presence of a resident soil microbiome (Schmitz et al., 2022), and increased wheat biomass under water stress when members were selected for osmotolerance (Agoussar et al., 2025). Such microbial consortia, including both bacteria and fungi, may provide broader functional coverage, with different members contributing to soil aggregation, metabolite production, nutrient exchange, and access to water beyond rhizosphere depletion zones (Liu et al., 2024). As a result, these complex multi-species consortia may improve plant-available water and enhance stress resilience at the system level, particularly under challenging climatic conditions and in degraded soils. However, most defined community studies have been conducted in gnotobiotic or pot systems and replicated evaluations of complex bacterial–fungal consortia in commercially relevant crops under deficit irrigation remain scarce.

To capture broad functional diversity, we assembled microbial consortia from a genome-sequenced culture collection of soil microorganisms, prioritizing spore-forming bacteria (with emphasis on *Streptomyces* and Bacillales), complemented by non-spore-forming Actinomycetota and, in four of five consortia, arbuscular mycorrhizal fungi (AMF) and/or ectomycorrhizal fungi (EMF) (Fig. 1; Table 1). The five consortia (A–E) comprised 12–55 bacterial strains (11–33 species) and 0–11 fungal species, and differed in richness, in the relative dominance of *Streptomyces* versus Bacillales, and in fungal composition, while sharing a subset of core taxa. The aim of this study was to test whether these consortia mitigate the effects of deficit irrigation on soil-grown leafy greens. Specifically, we asked (i) whether consortium inoculation reduces the yield, harvest-timing and physiological penalties of a 30% irrigation deficit in lettuce and spinach, (ii) whether responses depend on application rate, and (iii) whether consortia of different composition differ in efficacy. Because the consortia comprise taxa with documented plant-growth-promoting and stress-mitigating activity, we hypothesized that (H1) inoculated plants under deficit irrigation would outperform untreated deficit-irrigated plants, (H2) the higher application rate would produce larger effects, and (H3) responses would differ among consortia and between crops. We tested these hypotheses in four replicated greenhouse trials, one lettuce and one spinach trial at each of two sites, under soil-based cultivation in the Valencia region of Spain.

**Figure 1.**
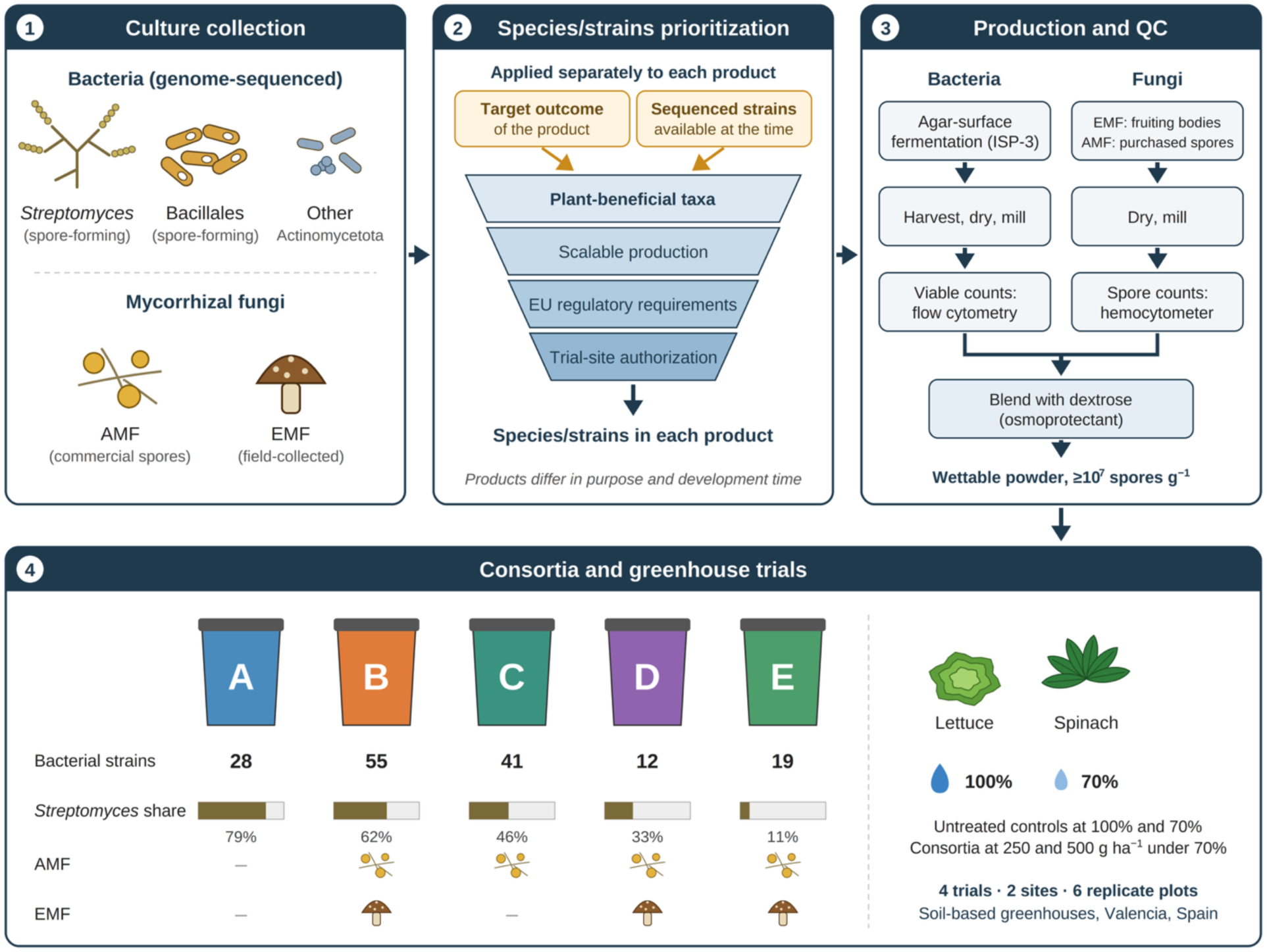
Composition and production of the microbial consortia. (1) Bacterial strains were drawn from a genome-sequenced culture collection dominated by spore-forming *Streptomyces* and Bacillales, with a smaller number of other Actinomycetota; mycorrhizal fungi comprised commercially sourced arbuscular mycorrhizal fungi (AMF) and field-collected ectomycorrhizal fungi (EMF). (2) For each product separately, species and strains were prioritized according to the product’s target outcome and the sequenced strains available at the time of its development, from taxa with documented plant-beneficial activity, and further constrained by scalable production, and EU regulatory requirements. (3) Bacteria were produced by agar-surface fermentation, harvested, dried and milled, with viable counts determined by flow cytometry; fungal spores were dried and milled, with spore counts determined by hemocytometer. Components were blended with dextrose as an osmoprotectant into wettable powders (≥10⁷ spores g⁻¹). (4) The five consortia (A–E) differ in the number of bacterial strains, the share of *Streptomyces* among bacterial strains, and the presence of AMF and EMF (Table 1; Table S1); consortium colours match those in Figs 2–5. Consortia were tested at 250 and 500 g ha⁻¹ under 70% irrigation, alongside untreated controls at 100% and 70% irrigation, in four soil-based greenhouse trials (lettuce and spinach at each of two sites) in the Valencia region, Spain. Alt text: Four-panel flow diagram: a culture collection of bacteria and mycorrhizal fungi; a funnel of four species and strain prioritization criteria applied separately to each product, fed by its target outcome and the strains available at the time; parallel bacterial and fungal production and quality-control steps; and five coloured product packets labelled A to E showing their number of bacterial strains, Streptomyces share and fungal content, next to lettuce and spinach grown under 100% and 70% irrigation.

**Table 1.** Composition of microbial consortia A–E, given as the number of strains per species (bacteria) or the presence of each species (fungi). Designations such as ‘sp. AK04’ refer to the closest GTDB reference genome; ‘*Streptomyces* sp.’ denotes strains without a validly published species assignment (PCT patent publication WO2025097154, Microbial Compositions and Methods). Fungal entries represent spore inocula of each species rather than characterized strains: AMF were obtained from commercial suppliers and EMF spores from field-collected fruiting bodies. Strain-level membership is given in Table S1.

| Species | A | B | C | D | E |
|---|---|---|---|---|---|
| **Number of bacterial strains** | **28** | **55** | **41** | **12** | **19** |
| *Arthrobacter* sp. AK04 |  | 1 | 1 | 2 | 1 |
| *Arthrobacter* sp. StoSoilA2 |  | 1 | 1 | 1 |  |
| *Bacillus haynesii* |  | 1 | 1 | 1 | 1 |
| *Bacillus inaquosorum* |  | 2 | 1 | 1 | 1 |
| *Bacillus licheniformis* |  | 1 | 1 |  | 1 |
| *Bacillus paralicheniformis* |  |  |  |  | 1 |
| *Bacillus safensis* |  | 2 | 2 |  | 1 |
| *Bacillus spizizenii* |  | 1 |  |  | 1 |
| *Bacillus subtilis* |  | 2 | 3 |  | 1 |
| *Bacillus velezensis* | 1 |  | 2 |  |  |
| *Kocuria oxytropis* |  | 1 |  |  | 1 |
| *Lysinibacillus capsici* |  | 1 |  | 1 |  |
| *Lysinibacillus sphaericus* | 1 |  |  |  |  |
| *Microbacterium* sp. MYb45 | 1 | 1 | 1 |  |  |
| *Paenibacillus cellulositrophicus* | 2 | 1 | 2 | 1 | 2 |
| *Peribacillus frigoritolerans* |  | 2 | 2 |  | 3 |
| *Priestia aryabhattai* |  | 1 | 2 | 1 | 1 |
| *Priestia endophytica* |  | 1 | 1 |  | 1 |
| *Priestia megaterium* |  | 1 | 1 |  | 1 |
| *Rhodococcus baikonurensis* | 1 | 1 | 1 |  |  |
| *Streptomyces albogriseolus* |  | 1 | 1 |  |  |
| *Streptomyces bacillaris* | 2 | 2 | 1 |  |  |
| *Streptomyces bikiniensis* |  | 2 |  |  |  |
| *Streptomyces eurythermus* |  | 1 |  |  |  |
| *Streptomyces fimbriatus* |  | 1 | 1 |  |  |
| *Streptomyces globosus* | 4 | 7 | 5 | 1 |  |
| *Streptomyces griseoaurantiacus* |  | 1 | 1 |  |  |
| *Streptomyces griseoincarnatus* | 2 | 1 |  |  |  |
| *Streptomyces griseoviridis* |  | 1 |  |  | 1 |
| *Streptomyces misionensis* | 2 | 6 | 5 | 1 |  |
| *Streptomyces murinus* | 3 | 1 | 1 |  |  |
| *Streptomyces nogalater* | 1 |  |  |  |  |
| *Streptomyces olivaceus* |  | 1 |  | 1 |  |
| *Streptomyces rameus* | 2 | 2 |  | 1 |  |
| *Streptomyces rochei* |  | 2 | 4 |  |  |
| *Streptomyces sennicomposti* | 3 | 3 |  |  |  |
| *Streptomyces* sp. | 3 | 2 |  |  | 1 |
| **Number of fungal species** | **0** | **11** | **4** | **8** | **11** |
| *Claroideoglomus etunicatum* |  | 1 | 1 | 1 | 1 |
| *Funneliformis mosseae* |  | 1 | 1 | 1 | 1 |
| *Glomus aggregatum* |  | 1 | 1 | 1 | 1 |
| *Pisolithus tinctorius* |  | 1 |  |  | 1 |
| *Rhizophagus irregularis* |  | 1 | 1 | 1 | 1 |
| *Rhizopogon amylopogon* |  | 1 |  | 1 | 1 |
| *Rhizopogon fulvigleba* |  | 1 |  | 1 | 1 |
| *Rhizopogon luteolus* |  | 1 |  | 1 | 1 |
| *Rhizopogon villosulus* |  | 1 |  | 1 | 1 |
| *Scleroderma cepa* |  | 1 |  |  | 1 |
| *Scleroderma citrinum* |  | 1 |  |  | 1 |

## Materials and methods

### Strain origin, selection and consortium design

All bacterial strains were drawn from the Oath Inc. culture collection, which comprises isolates obtained from soils in the USA and Australia. Each consortium was originally developed to include taxa with documented plant-beneficial activity, with an emphasis on spore-forming *Streptomyces* and Bacillales (*Bacillus*, *Lysinibacillus*, *Paenibacillus*, *Peribacillus* and *Priestia*), because spores tolerate drying, formulation and storage and allow reproducible viable counts in a wettable-powder product (Bashan et al., 2014), and because *Streptomyces* persist in soil as spores and are known for rhizosphere and endophytic colonization and plant-growth promotion (Viaene et al., 2016; Worsley et al., 2020). A smaller number of non-spore-forming Actinomycetota (*Arthrobacter*, *Kocuria*, and *Rhodococcus*) were also included for their plant-growth-promoting potential. Strains of these genera have been reported to promote root growth and nutrient acquisition. ACC deaminase-containing *Rhodococcus* strains stimulated root elongation of Indian mustard (Belimov et al., 2005), and ACC deaminase-containing *Arthrobacter protophormiae* improved nodulation and mycorrhization of pea (Barnawal et al., 2014). *Kocuria rhizophila* solubilizes phosphate, produces indole-3-acetic acid and promotes growth of maize and tomato (Li et al., 2020; Mauceri et al., 2024). These genera are also desiccation-tolerant members of soil communities (LeBlanc et al., 2008). Strain selection also aimed to include both motile and non-motile taxa, as motility is a key determinant of whether an introduced strain disperses through the soil pore network, reaches the rhizosphere, and competes for colonization sites on developing roots (Wang et al. 2026).

All strains were subjected to a genome-based safety screen. Assembled genomes were searched for antimicrobial resistance (AMR) determinants against two curated databases, CARD (Alcock et al., 2023) and ResFinder (Bortolaia et al., 2020), and for virulence factors against VFDB (Liu et al., 2022). Following the European Food Safety Authority (EFSA) guidance on whole-genome sequence analysis of microorganisms intentionally used in the food chain, applied here as a conservative and widely recognized standard although these products are not placed on the market as food-chain microorganisms, hits with a minimum of 80% identity and 70% length coverage of the subject sequence were reported and evaluated (EFSA, 2024). Reported hits were assessed for genomic context to distinguish chromosomally encoded (intrinsic) determinants from plasmid-borne or otherwise mobile (acquired) determinants, since only acquired resistance to clinically relevant antimicrobials is considered a safety concern. Genomes were additionally analyzed with antiSMASH (Blin et al., 2023) to identify biosynthetic gene clusters with the predicted capacity to produce antimicrobials classified as medically important for human medicine (WHO, 2024). No strain in consortia A–E carried acquired AMR determinants against clinically relevant antimicrobials, virulence factors of concern, or the predicted capacity to produce medically important antimicrobials; all strains passed the screen, and none was excluded on this basis. Literature searches found no associations between the selected species and plants or animal disease.

Final compositions were further constrained by (i) the ability to produce each strain at relevant scale using existing fermentation protocols, and (ii) regulatory requirements for microbial products in the EU. In some cases, strains were replaced during product development, for example when an originally selected strain met the exclusion criterion above or failed to grow at adequate rates. Because lettuce and spinach are not ectomycorrhizal hosts, and spinach (Amaranthaceae) is generally considered non-mycorrhizal (Brundrett & Tedersoo, 2018), the fungal components were not expected to form functional symbioses with both crops in these trials; they were retained because the consortia were also developed as products for use across other cropping systems.

The five consortia differ in richness (12–55 bacterial strains), in the proportion of *Streptomyces* (79% of bacterial strains in consortium A, 62% in B, 46% in C, 33% in D and 11% in E) and in fungal composition (none in A; AMF only in C; AMF plus EMF in B, D and E). These differences reflect the fact that each consortium was developed for a different target outcome and at a different time, when different numbers of sequenced strains were available in the collection, together with the constraints described above. They were not systematically varied design factors, and the study was not designed to isolate the effect of any single compositional variable. Several taxa were shared across consortia, including *Paenibacillus cellulositrophicus* in all five and *Streptomyces globosus* and *S. misionensis* in consortia A–D (Table 1; Table S1). The consortia correspond to compositions described in patent publications WO2025097154 and WO2025240938.

EMF fruiting bodies were collected in the United States, which is not a Party to the Convention on Biological Diversity or to its Nagoya Protocol; access and benefit-sharing obligations under the Protocol and under Regulation (EU) No 511/2014 therefore do not apply to these genetic resources. AMF inocula were purchased from commercial suppliers in Europe.

### Taxonomic classification of bacteria and fungi

Genomic sequences used for the identification of bacterial strains (all derived from Oath Inc.’s strain collection) were obtained through whole-genome sequencing of the strains listed in Table 1 and Table S1, using either Illumina paired-end or Oxford Nanopore sequencing. Illumina paired-end reads were preprocessed using Fastp (Chen et al., 2018) and preprocessed reads were assembled using SPAdes (Bankevich et al., 2012) through VEBA (Espinoza & Dupont, 2022; Espinoza et al., 2024). Oxford Nanopore reads were preprocessed using Fastplong (Chen, 2023), preprocessed reads were assembled with Flye (Kolmogorov et al., 2019), and assemblies were polished using Medaka implemented through VEBA. Each assembly met high-quality genome standards (Bowers et al., 2017) as assessed with CheckM2 (Chklovski et al., 2023), and taxonomy was classified using GTDB-Tk (release 226) (Chaumeil et al., 2022). Arbuscular mycorrhiza fungal spores from *Rhizophagus irregularis, Claroideoglomus etunicatum* and *Funneliformis mosseae* were classified by the supplier Mycointech, Tarragona, Spain. *Glomus aggregatum* was characterized by the supplier UmaHari LLC, New York, USA. Ectomycorrhiza fungi spores were obtained from *Pisolithus tinctorius, Rhizopogon amylopogon, Rhizopogon fulvigleba, Rhizopogon luteolus, Rhizopogon villosulus*, *Scleroderma cepa* and *Scleroderma citrinum* by collecting whole fruiting bodies in the southern Oregon Cascades, the United States during 2024 and 2025 by Dr. Michael Amaranthus at Mycorrhizal Applications Inc., OR, USA. Fruiting body morphological characteristics for each species were carefully examined in the field. In the laboratory, spores were microscopically examined for size, shape and color. Spore counts were quantified with a hemocytometer. To process the fruiting bodies the sterile bases of whole fruiting bodies were removed, and the remaining spore bearing tissue cleaned and sliced for drying. The spore-containing tissues were dried at 54°C for 100 h in a dehydrator and ground into a fine powder with a steel food grade grinder. Finished concentrated powders were tagged and sealed in a sterile plastic bag for each specimen.

### Microbial consortium formulation

Bacterial components of products A–E were produced via agar surface fermentation on ISP Medium No. 3 (HiMedia M358). Working stocks were revived from −80 °C glycerol stocks into glucose broth seed cultures, verified for purity by single-colony morphology on ISP-3 plates, and scaled to production plates incubated at 28 °C until visible sporulation. Biomass was harvested by scraping, blended with dextrose as an osmoprotectant, dried at ambient conditions, and milled to a homogeneous wettable powder. Live cell and spore count for each bacterial strain were quantified by flow cytometry (ISO 19344 Part B; BioForm Solutions Inc., San Diego, CA). EMF spores were sourced from field-collected fruiting bodies harvested in the southern Oregon Cascades (USA) and processed by Mycorrhizal Applications Inc. (OR, USA); AMF spores were obtained commercially (Mycointech, Tarragona, Spain; UmaHari LLC, New York, USA). Bacterial strains were blended in equal proportions based on flow cytometry counts and bacterial and fungal components were combined with dextrose to a final count of ≥10⁷ spores g⁻¹ and packaged as wettable powder.

### Trial sites, growing conditions and experimental design

All trials were conducted by the test facility GMW Bioscience (Alberic, Valencia, Spain) under Good Experimental Practice (GEP accreditation EOR67/08) and followed EPPO PP 1/135(4), PP 1/152(4) and PP 1/181(5) (EPPO 2012, 2014, 2021) and CEN 17700-1 and CEN 17700-3. Four trials were conducted in spring 2025 at two sites, with lettuce (*Lactuca sativa*, variety Romana Cercas) and spinach (*Spinacia oleracea*, variety Grebe RZ) at Torrent and at Alberic, Valencia (Fig. 2B; Table S2). Site, soil, management, application and harvest parameters for each trial are given in Table S2; the text below describes features common to all four trials and the between-site differences that bear on interpretation.

**Figure 2.**
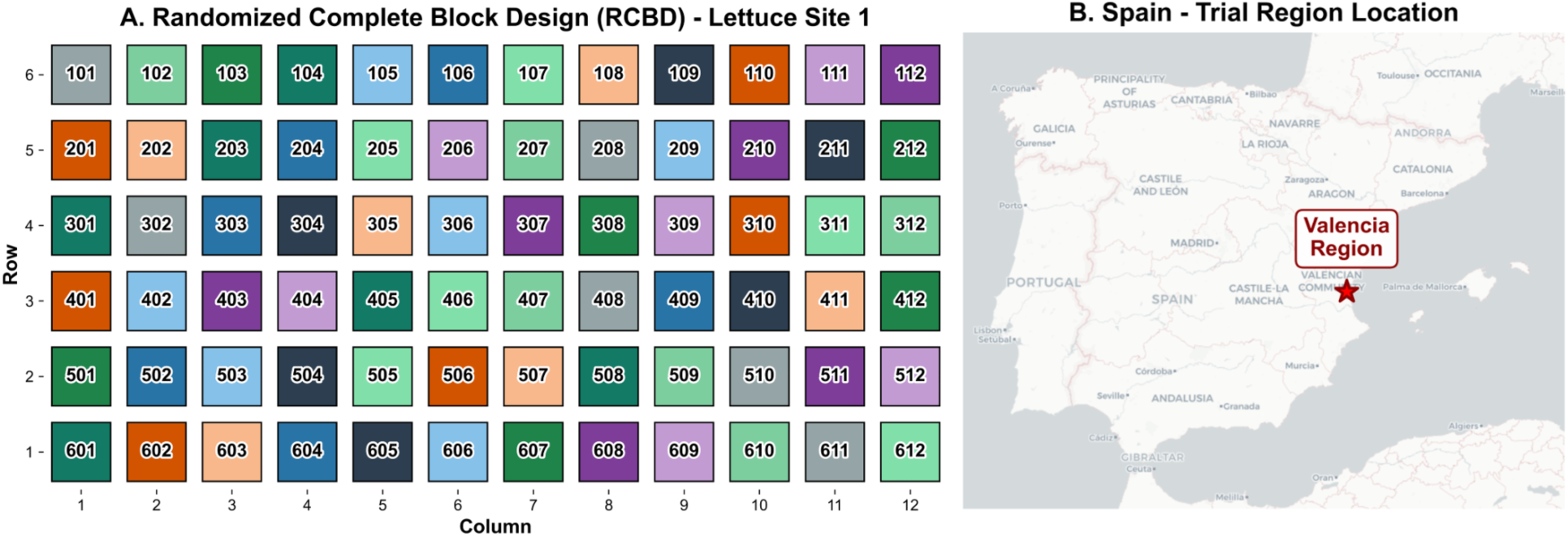
Trial location and experimental design. (A) Randomized complete block design (RCBD) layout of trial 009 (lettuce, Torrent): 72 plots arranged in six blocks (rows), with the 12 treatments randomized within each block. Treatments were untreated controls at 100% (UTC 100%; dark slate) and 70% irrigation (UTC 70%; grey), and consortia A (blue), B (orange), C (teal), D (purple) and E (green), each applied at a low (250 g ha⁻¹; light shade) and a high (500 g ha⁻¹; dark shade) rate under 70% irrigation. Layouts of the other three trials followed the same design with site-specific plot dimensions and spacing (Table S2). (B) Location of the trial region (Valencia, Spain). Details of the four trials are given in Table S2. Alt text: Left, a grid of 72 numbered, colour-coded plots in six rows and twelve columns showing the randomized treatment layout. Right, a map of Spain with a star marking the Valencia region.

Plants were grown in soil-based cultivation in the local soil of each greenhouse; no substrate or soilless growing medium was used. All four soils fell in the clay textural class but differed markedly between sites, with 50–54% clay and pH 6.9–7.1 at Torrent against 79–80% clay and pH 7.4–8.3 at Alberic (Table S2).

Trials were conducted in multi-span plastic-covered greenhouses with passive ventilation. Air temperature and relative humidity inside each greenhouse were logged as daily minima and maxima only, so daily means are not available; summary values are given in Table S2. Conditions were consistently hotter and drier at Alberic, where daily maxima reached ≥40 °C on 27–32 of the trial days and peaked at 61.3 °C, against 17–19 days and 53.2 °C at Torrent.

Seedlings were transplanted by hand between 5 and 14 March 2025 at 6.06 plants m⁻² (Torrent) or 4.80 plants m⁻² (Alberic); planting density, row spacing and plot dimensions were set by site and were identical for the two crops within a site (Table S2). Fertilization differed by site and is not comparable across trials: both Torrent trials received a single pre-planting base dressing of NPK 15-15-15 and a sheep-pellet organic amendment with no fertigation, whereas both Alberic trials were fertigated three times through the drip system, including a substantial nitrogen and potassium input on 26 March 2025 during the stress period (Table S2).

Each trial comprised 12 treatments with six replicate plots each (72 plots per trial; plots of approximately 10 m²; Fig. 2A; Table S2): untreated controls at 100% (UTC 100%) and 70% (UTC 70%) irrigation, and consortia A–E each applied at a low (250 g ha⁻¹) and a high (500 g ha⁻¹) rate under 70% irrigation. Plots were arranged in six rows of 12, and each row contained one plot of each treatment in randomized order, giving a randomized complete block layout with rows as blocks. Because blocking did not target a specific environmental gradient, spatial heterogeneity was accounted for in the analysis by Gaussian-process detrending on plot coordinates rather than by a block term (see Statistical analysis). Each plot comprised four crop rows and had an independent irrigation supply, so that 100% and 70% plots could be irrigated independently; adjacent plots were separated by 80–100 cm. To minimize edge effects and lateral water movement between neighbouring plots with different irrigation regimes, only the two central rows of each plot were assessed and harvested. The fully irrigated control received 100% of calculated crop water requirements, whereas the deficit irrigation treatments received 70%. Soil water content was not measured.

### Microbial inoculant application protocol

All plots received uniform irrigation for the first four to eight days after transplanting to support crop establishment. Irrigation was then differentiated by treatment and maintained until harvest, with deficit applied to the inoculant-treated plots and to the 70% irrigation control. Irrigation was delivered by drip, with one 2 L h⁻¹ emitter per plant on six days per week, and the regime was set by site rather than by crop: at Torrent, 45 min per irrigation day under full irrigation and 30 min under deficit (1.50 and 1.00 L plant⁻¹); at Alberic, 40 and 25 min (1.33 and 0.83 L plant⁻¹) (Table S2). The deficit treatments are designated ’70% irrigation’ throughout, following the trial protocol, but the volumes actually delivered corresponded to 67% of the fully irrigated control at Torrent and 62.5% at Alberic. At Alberic, the drip system was also used for fertigation (Table S2). The first fertigation (13 March) preceded the onset of deficit irrigation, but the second and third (26 March and 9 April) were applied after irrigation regimes had been differentiated. Whether fertilizer was dosed per plot or injected at a fixed concentration into the irrigation water was not recorded; in the latter case, deficit-irrigated plots would have received approximately 62.5% of the N and K delivered to the fully irrigated control on those dates. At Torrent, all fertilizer was applied before transplanting and was uniform across plots.

Microbial consortia were suspended in water at 0.109 and 0.217 g L⁻¹ (Torrent) or 0.100 and 0.200 g L⁻¹ (Alberic) for the low and high rates respectively, and applied as soil-directed treatments using a knapsack sprayer fitted with a single SJ303 drench nozzle. Untreated control plots received neither inoculant nor compensatory water. Application volume was 2300 L ha⁻¹ at Torrent and 2500 L ha⁻¹ at Alberic, equivalent to 38 and 52 mL per plant at each of three application events. The first application was made within 24 h of transplanting and the second 14 days later in all four trials; the third followed 14 days after the second except in trial 009, where it was made after 18 days (Table S2). Crop stage at the third application ranged from BBCH 19 to BBCH 43 across trials. Application rates were 250 g ha⁻¹ (low) and 500 g ha⁻¹ (high) of formulated product per application corresponding to 2.5 × 10⁹ and 5.0 × 10⁹ CFU ha⁻¹, or 4.1 × 10⁴ and 8.3 × 10⁴ CFU plant⁻¹ at Torrent and 5.2 × 10⁴ and 1.0 × 10⁵ CFU plant⁻¹ at Alberic. All inoculant applications were timed to coincide with early crop establishment and subsequent developmental stages to maximize potential effects on root colonization, rhizosphere function, and plant performance under water-limited conditions. All inoculant formulations included 0.1% v/v Tween 80 as a wetting agent to aid dispersion of the inoculant in the spray suspension; the test facility nevertheless reported difficulty in diluting the product and observed flocculation in the tank mix in all four trials, and a specialized nozzle was used to achieve even distribution.

### Plant phenotypic measurements and harvest

Measurements followed EPPO guidance and were made on plants in the two central rows of each plot at the last assessment before harvest (BBCH ≥ 48; Lancashire et al., 1991). Sample sizes differed by variable: 12 leaves per plot for leaf reflectance indices, chlorophyll index and leaf area; six leaves per plot for leaf length; six plants per plot for fresh and dry aerial and root biomass; six roots per plot for root length; and 12 plants per plot for individual plant weight and head diameter. Phytotoxicity, crop vigor, growth stage, canopy cover and wilting were scored at whole-plot level.

Leaf chlorophyll content was estimated with a as a chlorophyll index derived from leaf reflectance, recorded with a CI-710S leaf spectrometer (CID Bio-Science, Camas, WA, USA) on 12 leaves per plot. (Markwell et al., 1995). Leaf reflectance spectra were recorded on the same leaves with a CI-710S leaf spectrometer (CID Bio-Science, Camas, WA, USA), and used to derive a water band index (WBI), computed by the instrument software from the ratio of reflectance at 970 nm to that at 900 nm, so that higher values indicate greater water stress and lower leaf water content (Peñuelas et al., 1993). Wilting incidence was scored visually at each assessment as the percentage severity rating for the plot as a whole. Root length (m) was measured on six carefully excavated and washed roots per plot. Yield was determined as the fresh marketable weight of the plants harvested from a defined area within the two central rows of each plot, 1.98–2.49 m², corresponding to 12 plants in every trial (Table S2), and expressed per hectare.

All plots within a trial were harvested on the same day, before the onset of deterioration in the most advanced plots; at that time some plants had not yet reached their optimal developmental stage. For each plot, the assessor estimated the number of additional days required for plants to reach optimal harvest stage; this value is reported as harvest delay (days), with 0 indicating that plants were at optimal stage on the harvest date. Estimated delays for the stressed untreated control ranged from 3 to 6 days across trials (Table S2).

### Statistical analysis

Analyses were performed for lettuce and spinach across two sites per crop, with plot-level measurements as the unit of analysis. Harvest-stage observations were identified using BBCH thresholds (Lancashire et al., 1991) and analyzed at the last harvest-relevant assessment date per site (BBCH ≥ 48). For continuous physiological traits and yield, spatial variation within each site was adjusted before treatment modeling using Gaussian-process (GP) regression on plot coordinates (Gilmour et al., 1997; Rodríguez-Álvarez et al., 2018). A fixed radial basis function spatial trend (ConstantKernel × RBF, length-scale fixed at 5.0 with alpha = 0.01, selected empirically by comparing spatial trend surfaces and detrended residual maps) was fitted to plot coordinates, and detrended values were computed as residualized trait values centered on the site mean. Spatial detrending was performed as a preliminary step prior to mixed-effects modeling rather than through joint modeling of spatial structure and treatment effects in a single model. Joint modeling, as in the AR1×AR1 framework of Gilmour et al. (1997), offers the advantage of propagating spatial uncertainty directly into treatment inference, but assumes spatial correlation is separable across rows and columns independently. The GP with a RBF kernel captures non-separable two-dimensional smooth surfaces, accommodating gradients that do not align with plot axes. This two-step approach is a pragmatic trade-off prioritizing richer spatial modeling; the consistency of results with independent mixed-model re-analysis confirms this does not materially affect treatment inference (Pérez-Valencia et al., 2022). Across-site treatment effects were then estimated with linear mixed-effects models (Laird & Ware, 1982; Pinheiro & Bates, 2009), with treatment as a fixed effect and site as a random intercept. Treatment contrasts were evaluated against two untreated controls: UTC at 70% irrigation (primary drought-control comparison) and UTC at 100% irrigation (well-watered reference). Wald-type z-tests were used for fixed-effect inference (McCullagh, 2019; McCullagh & Nelder, 1989). Harvest delay (days) was treated as a discrete endpoint and analyzed using generalized estimating equations (Poisson family, exchangeable working correlation) with site-level clustering (Liang & Zeger, 1986); treatment means are reported as back-transformed expected values on the response scale. P-values for treatment contrasts were corrected for multiple testing using the Benjamini-Hochberg false discovery-rate procedure (Benjamini & Hochberg, 1995), applied separately for the UTC 70% and UTC 100% contrast families. Reported uncertainty bars correspond to model-based standard errors of estimated treatment means. Analyses were run in Python using pandas (McKinney, 2010), numpy (Harris et al., 2020), statsmodels (Seabold & Perktold, 2010), scikit-learn (Pedregosa et al., 2011), and scipy (Virtanen et al., 2020).

## Results

### Microbial consortia improve yield under deficit irrigation in leafy greens

Deficit irrigation reduced the yield of untreated plants by approximately 10% in lettuce and 13% in spinach (UTC 70% vs UTC 100%, P < 0.001; Fig. 3). Most consortium treatments did not fully compensate for this loss: under deficit irrigation, yields in 8 of 10 lettuce and 7 of 10 spinach treatments remained significantly lower than in UTC 100% (P < 0.05). Consortium inoculation nevertheless increased yield relative to UTC 70%, by approximately 2–8% in lettuce (0.9–3.0 t ha⁻¹) and 4–12% in spinach (1.1–3.3 t ha⁻¹) depending on consortium and application rate. These increases were statistically significant for 5 of 10 lettuce treatments (consortia B, C, D and E at the high rate and C at the low rate) and 6 of 10 spinach treatments (A at both rates, B and C at the high rate, and E at both rates; P < 0.05; Fig. 3, Fig. S1).

**Figure 3.**
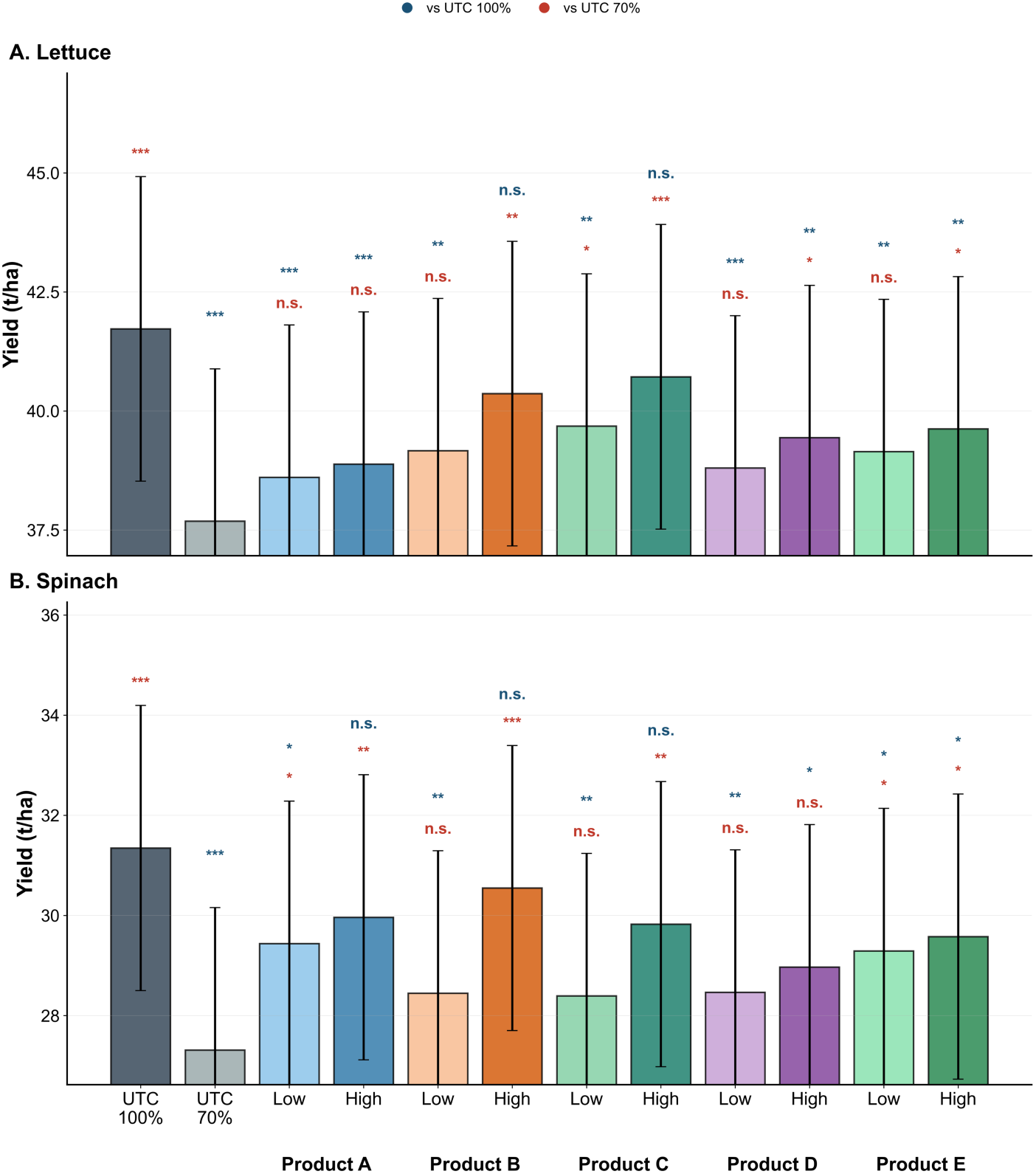
Yield of (A) lettuce and (B) spinach under full and deficit irrigation. Treatments were untreated controls at 100% (UTC 100%) and 70% irrigation (UTC 70%), and consortia A–E (labelled Product A–E) applied at a low (250 g ha^−1^) or high (500 g ha^−1^) rate under 70% irrigation. Bars show spatially detrended, model-estimated treatment means across two sites per crop (linear mixed-effects model with site as a random intercept); error bars show model-based standard errors of the means, which include between-site variance, so overlapping error bars do not imply non-significant contrasts. Significance labels are stacked above each bar: the upper label (dark blue) gives the comparison with UTC 100% and the lower label (red) the comparison with UTC 70% (Benjamini–Hochberg-adjusted Wald tests; ***P < 0.001, **P < 0.01, *P < 0.05; n.s., not significant). The y-axes do not start at zero. Plot-level data for each site are shown in Fig. S1. Alt text: Two bar charts of yield for lettuce (top) and spinach (bottom). The 70% irrigation control yields less than the fully irrigated control; the ten consortium treatments are intermediate, and high-rate treatments generally yield more than low-rate treatments.

For every consortium in both crops, the high application rate (500 g ha^−1^) produced a numerically greater yield than the low rate (250 g ha^−1^), but these differences were not significant (P > 0.05). The five consortia (A–E) differed in the magnitude of their response. In lettuce, consortium C gave the largest yield gains, and at the high rate consortia B and C were the only treatments with yields not significantly different from UTC 100% (P > 0.05); consortium A showed the smallest improvement and did not differ significantly from UTC 70% at either rate. In spinach, the high rate of consortia A, B and C produced yields not significantly different from UTC 100% (P > 0.05), whereas consortium D did not differ significantly from UTC 70% at either rate (Fig. 3). Because absence of a significant difference does not demonstrate equivalence, these findings indicate that microbial inoculation partially offset, but in most cases did not eliminate, the yield penalty of deficit irrigation.

### Consortium inoculation reduced harvest delay under deficit irrigation

In untreated plots, deficit irrigation delayed the optimal harvest stage by approximately 4.9 days in lettuce and 4.4 days in spinach, compared with 0 and 0.4 days, respectively, under full irrigation (Fig. 4). In lettuce, all consortium treatments significantly reduced harvest delay relative to UTC 70%, by approximately 2.9–4.7 days (P < 0.01), but all remained significantly delayed relative to UTC 100%, which showed no delay (P < 0.001). In spinach, nine of ten treatments significantly reduced harvest delay relative to UTC 70%, by approximately 2.4–4.4 days (P < 0.01); the exception was consortium B at the low rate, which was highly variable and did not differ significantly from either control. Five spinach treatments (A and B at the high rate, D at both rates, and E at the low rate) did not differ significantly from UTC 100%, whereas consortia A and C at the low rate remained significantly more delayed than UTC 100%. Consortia C and E at the high rate showed no harvest delay in any plot and were therefore significantly less delayed than UTC 100%; because these estimates lie on the boundary of the model’s parameter space, these contrasts should be interpreted with caution. The high rate resulted in a shorter harvest delay than the low rate for all five consortia in lettuce and for four of five in spinach (all except D).

**Figure 4.**
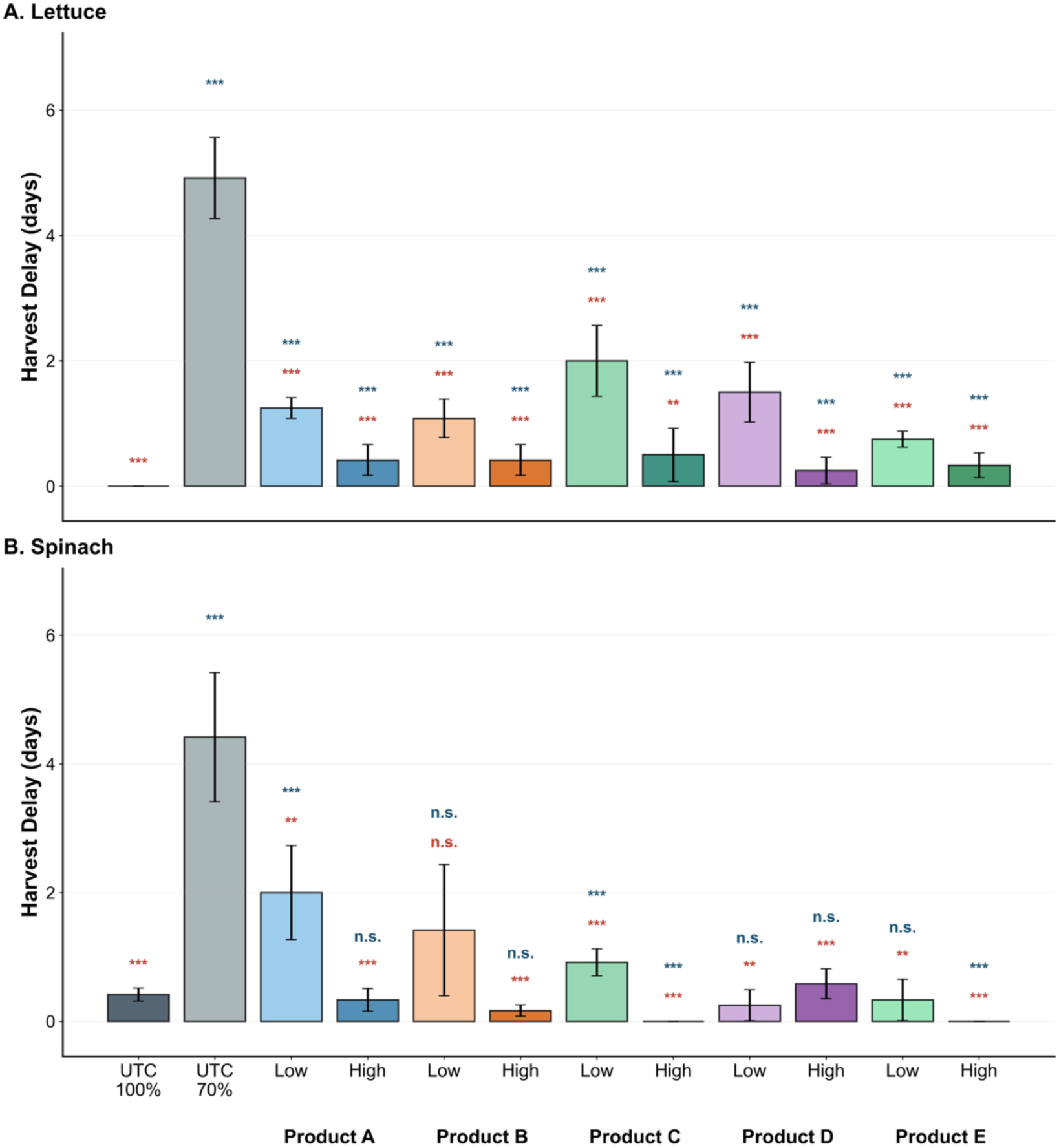
Harvest delay of (A) lettuce and (B) spinach under full and deficit irrigation. Harvest delay is the estimated number of additional days required to reach optimal harvest stage on the common harvest date. Treatments as in Fig. 3. Bars show back-transformed means from generalized estimating equations (Poisson family, site-level clustering) ± model-based standard errors. Significance labels as in Fig. 3: upper (dark blue), comparison with UTC 100%; lower (red), comparison with UTC 70% (Benjamini–Hochberg-adjusted; ***P < 0.001, **P < 0.01, *P < 0.05; n.s., not significant). Bars without error bars are treatments with zero delay in all plots; contrasts involving these treatments should be interpreted with caution. Alt text: Two bar charts of harvest delay in days for lettuce and spinach. The 70% irrigation control shows a delay of about 4 to 5 days, most consortium treatments show delays of 0 to 2 days, and the fully irrigated control shows little or no delay.

### Consortium inoculation improved physiological status at harvest

Deficit irrigation reduced root length and increased wilting incidence and WBI in untreated plants of both crops, and reduced SPAD in spinach; in lettuce, the lower SPAD of UTC 70% was not significantly different from UTC 100% (Fig. 5). In lettuce, all treatments except consortium C at the low rate significantly increased root length relative to UTC 70%, by approximately 5–12% (P < 0.05), but only consortia D and E at the high rate were not significantly different from UTC 100%. In spinach, root length was significantly greater than in UTC 70% only for the high rate of consortia A, C, D and E (approximately 7–12%; P < 0.05), and only A and C at the high rate were not significantly different from UTC 100%. All consortium treatments in both crops had significantly lower WBI than UTC 70% (lettuce, P < 0.01; spinach, P < 0.05) and did not differ significantly from UTC 100%. In lettuce, SPAD was significantly higher than in UTC 70% only for consortium B at the high rate and consortium E at both rates (P < 0.05), and no treatment differed significantly from UTC 100%. In spinach, eight of ten treatments (all except B and D at the low rate) significantly increased SPAD relative to UTC 70% (P < 0.05), and five (A, B and C at the high rate, and E at both rates) did not differ significantly from UTC 100%. In both crops, all consortium treatments significantly reduced wilting incidence relative to UTC 70% (lettuce, from approximately 26% to 7–15%; spinach, from approximately 33% to 14–19%; P < 0.001), but wilting remained significantly higher than in UTC 100% (≤1%; P < 0.001).

**Figure 5.**
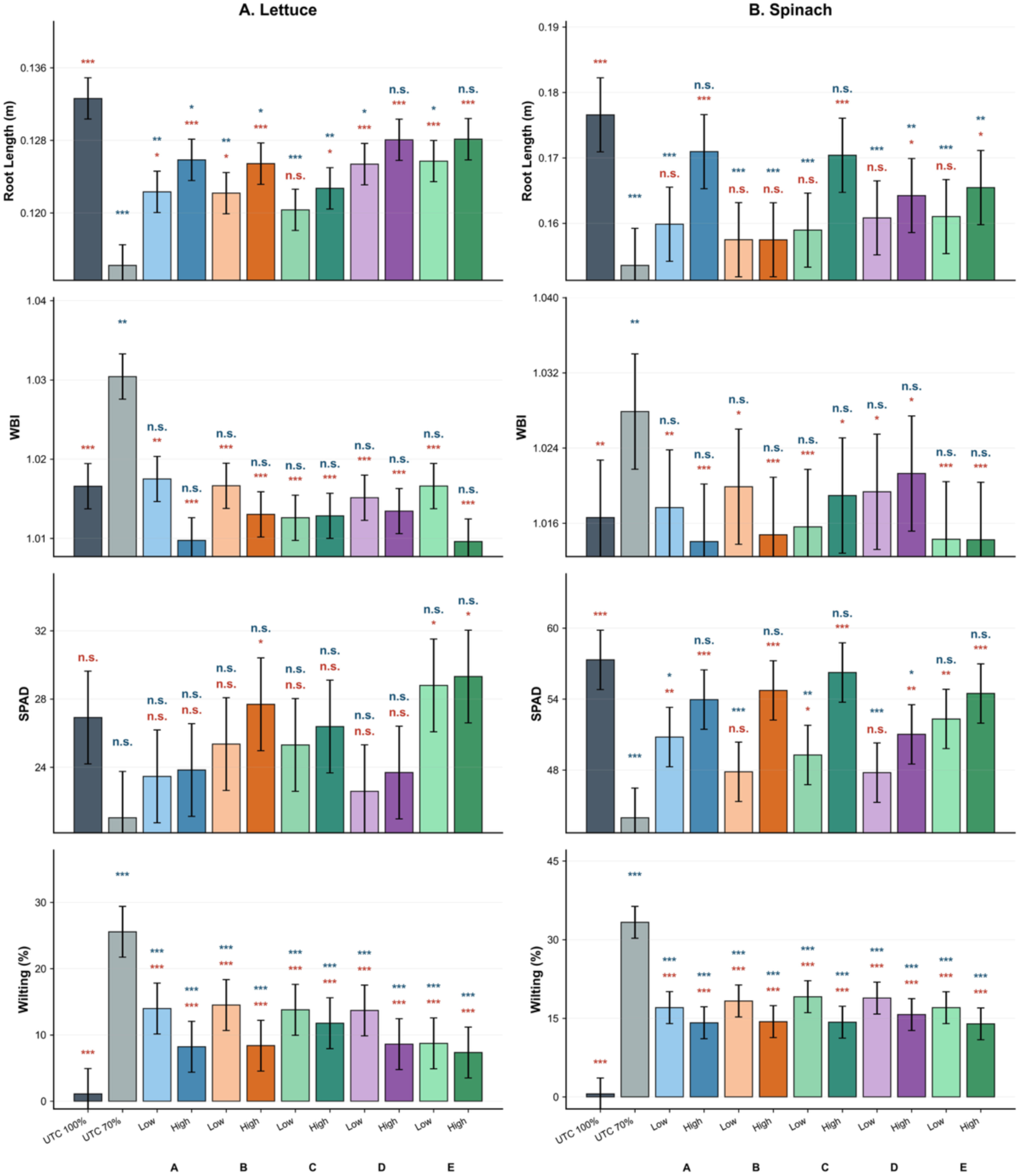
Physiological status at harvest of (A) lettuce and (B) spinach under full and deficit irrigation. Rows show root length (m), Water Band Index (WBI; dimensionless), leaf chlorophyll content (SPAD units) and wilting incidence (% of plants). Treatments as in Fig. 3. Bars show spatially detrended, model-estimated means ± model-based standard errors. Significance labels as in Fig. 3: upper (dark blue), comparison with UTC 100%; lower (red), comparison with UTC 70% (Benjamini–Hochberg-adjusted; ***P < 0.001, **P < 0.01, *P < 0.05; n.s., not significant). The y-axes for root length, WBI and SPAD do not start at zero. Alt text: Eight bar charts in four rows and two columns (lettuce left, spinach right) showing root length, Water Band Index, SPAD and wilting. The 70% irrigation control has the shortest roots, highest WBI, lowest SPAD and highest wilting; consortium treatments are generally intermediate between the two controls.

## Discussion

This study tested whether complex, multi-species microbial consortia mitigate the effects of a 30% irrigation deficit in soil-grown lettuce and spinach. Our first hypothesis (H1) was partially supported. Inoculated plants under deficit irrigation consistently showed lower wilting incidence and WBI and, in most cases, shorter harvest delay and longer roots than untreated deficit-irrigated plants, but yield increases were significant in only about half of the treatments, and most treatments did not restore yield to the level of fully irrigated plants. The second hypothesis (H2), that the higher application rate would produce larger effects, was supported as a consistent directional trend but not statistically. The third hypothesis (H3) was supported: responses differed among consortia and between crops. Together, these results indicate that microbial inoculation can reduce, but generally not eliminate, the penalties of water limitation in leafy greens.

One of the most practically relevant outcomes was the reduction in harvest delay. Across both crops, most microbial treatments reduced delay by approximately 2.4–4.7 days relative to untreated deficit-irrigated controls. This is important not only as an indicator of improved stress tolerance, but also because tighter and more predictable harvest windows have direct operational value for growers through improved labor planning, harvest efficiency, and crop uniformity (Lindemann-Zutz et al., 2016). In high-value leafy green production systems, even modest shifts in harvest timing can influence marketability and postharvest handling. Because harvest delay was estimated by assessors rather than measured from staggered harvests, confirming this effect with direct measurements of time to marketable size would strengthen this conclusion.

The physiological measurements are consistent with improved plant performance under water stress. Root length was greater in most inoculated treatments than in untreated deficit-irrigated controls, suggesting improved belowground development and potentially enhanced access to water and nutrients. Root growth promotion under drought is a well-documented effect of plant growth-promoting bacteria, including *Streptomyces* and *Bacillus* spp., and has been linked to bacterial modulation of phytohormone and ethylene signalling (Glick, 2014; Vurukonda et al., 2016; Worsley et al., 2020). Likewise, the lower Water Band Index values observed across consortium treatments indicate reduced plant water stress, while decreases in wilting and, in several cases, improved chlorophyll status point to better maintenance of plant function during water limitation (Qiao et al., 2024; Wang et al., 2025). The convergence of these physiological and agronomic responses supports the conclusion that the observed effects reflect an improvement in crop resilience under deficit irrigation, although the agronomic responses were more variable than the physiological ones.

Treatment effects were not uniform across consortia or between crops. Some consortia performed better in lettuce, whereas others were more effective in spinach, indicating that consortium composition likely interacts with crop-specific physiology and rhizosphere ecology. These differences may reflect variation in microbial establishment, compatibility with host plants, functional complementarity among consortium members, or differential performance under specific soil and irrigation conditions. The fact that higher application rates generally produced stronger outcomes further suggests that inoculum density may influence the consistency or magnitude of rhizosphere colonization and downstream plant benefit, even though the rate effect was not statistically significant in the present dataset.

All five consortia produced at least some benefit under deficit irrigation. The consortia were not randomly assembled, but neither were they designed as a factorial test of composition: all were drawn from a pre-selected pool of taxa with documented plant-beneficial activity, their compositions were shaped by each product’s target outcome and development history as well as by manufacturing, and regulatory constraints, and they shared core taxa, including *Paenibacillus cellulositrophicus* in all five consortia and *Streptomyces globosus* and *S. misionensis* in four. The consortia also include species for which strains have been shown to improve drought tolerance, including *Bacillus velezensis* in wheat (Abd El-Daim et al., 2019), *Streptomyces olivaceus* in wheat under greenhouse and field conditions (Yandigeri et al., 2012), members of the *Bacillus subtilis* group (Zhang et al., 2010) and *Priestia megaterium* (Marulanda et al., 2009). However, drought-mitigating activity is strain-specific, and we did not test whether the strains used here share these traits. Species with documented drought-mitigating strains were concentrated in consortia B, C and E, yet consortium A, which contained few such species and no fungi, was among the most effective treatments in spinach. Species identity alone therefore does not explain the differences among consortia. In a comparison of inoculum design strategies for wheat under water stress, only an inoculum assembled from isolates able to grow under high osmolarity increased biomass under drought (Agoussar et al., 2025), and communities designed around specific functional traits have produced predictable host phenotypes under nutrient and salt stress (Herrera Paredes et al., 2018; Schmitz et al., 2022), consistent with the view that trait-based selection underpins resilient defined inoculants (de Souza et al., 2020; Shayanthan et al., 2022). The broadly similar responses across our consortia may therefore reflect their shared, pre-selected pool and shared membership rather than consortium-specific composition. Because the study included no randomly assembled, single-strain or carrier-only (dextrose) controls, we cannot attribute the benefit to particular members; resolving this will require strain-resolved genomic and phenotypic screening, for example for osmotolerance, combined with leave-one-group-out consortia.

The comparison among consortia nevertheless offers some insight into the role of the fungal fraction. Consortium A, which contained no fungi, performed comparably to the fungus-containing consortia, and in spinach its high rate was among the most effective treatments for yield and root length. Because neither crop forms ectomycorrhizae and spinach is generally non-mycorrhizal (Brundrett & Tedersoo, 2018), the bacterial fraction is the more likely contributor to the responses observed here, at least in spinach. Fungal inoculants also face particular challenges of establishment and competition with resident mycorrhizal communities in the field (Hart et al., 2018).

Two central open questions are whether the inoculated strains persisted in soil and how inoculation affected the resident soil microbial community; a related question is whether the degree of establishment predicted plant responses. None of these were measured in the present trials. Introduced microorganisms often decline rapidly in soil owing to competition with resident communities, predation and abiotic filtering (Kaminsky et al., 2019), yet inoculants can alter resident communities even when their own populations decline (Mawarda et al., 2020), so plant responses need not require long-term persistence. The ability of *Streptomyces* and Bacillales to persist in soil as spores (Viaene et al., 2016) makes establishment of these groups plausible but untested. Quantifying persistence and resident-community responses is therefore a priority of ongoing work, in which field trials are sampled repeatedly for rhizosphere and bulk-soil metagenomics with strain-resolved tracking of consortium members, these data will also test whether inoculant abundance predicts plant responses.

The mechanisms underlying these responses were not directly tested here, but the consortium composition suggests plausible biological explanations. Consortium members belong to taxa associated with rhizosphere colonization, soil aggregation and nutrient mobilization. Such functions may improve plant-available water indirectly through enhanced soil structure and water retention, or directly through improved root access to water and nutrients. However, the present study was designed to evaluate performance outcomes rather than resolve mechanism. Future work should therefore link agronomic responses to direct measurements of microbial establishment, soil physical properties, soil water content, plant hydraulic traits, and molecular indicators of stress response. Integrating metagenomics, metatranscriptomics, metatranslatomics and metabolomics (Moyne et al., 2023), with soil biophysical measurements will be especially valuable for determining which functions are most predictive of improved irrigation deficit stress response. Next steps also include open-field trials across contrasting soils and seasons, and dose–response designs spanning a wider range of application rates.

Several limitations should be considered when interpreting these findings. First, the study comprised four greenhouse trials conducted over a single growing season at two sites in one region, so the durability and reproducibility of the response under broader field conditions remain to be established. Second, all trials used soil-based cultivation, and the results cannot be extrapolated to soilless production systems, in which root microbiomes and irrigation management differ (Thomas et al., 2023). Third, soil water content was not measured, so the difference in soil moisture between irrigation regimes is inferred from irrigation volumes; untreated controls were not compensated for the small volume of water applied with the inoculants (less than 0.5% of seasonal irrigation). In addition, at Alberic fertilizer was partly delivered through the drip system after irrigation regimes diverged, so deficit-irrigated plots may have received less N and K than the fully irrigated control; comparisons with UTC 100% at this site, and the across-site estimates of the deficit-irrigation yield penalty, may therefore partly reflect nutrient as well as water supply. Comparisons between inoculated treatments and UTC 70% are not affected, as these plots received the same irrigation and fertigation regime. Fourth, the design lacked carrier-only and randomly assembled consortium controls, and inoculant persistence, resident-community responses and water productivity per unit input were not measured. Finally, while the results consistently show improved crop performance under reduced irrigation, the composite physiological response likely integrates multiple interacting processes, including soil structure, microbial activity, host physiology, and environmental context.

Despite these limitations, our results indicate that the introduction of complex microbial consortia can partially mitigate the effects of water limitation in leafy green production systems. By improving yield retention, reducing harvest delay, and moderating physiological stress under deficit irrigation, these consortia show promise as biologically based tools for climate-resilient agriculture. More broadly, the findings support the idea that functionally diverse microbial inoculants can become a component of resource-efficient crop management strategies as water scarcity intensifies across agricultural regions.

In summary, complex microbial consortia improved the performance of lettuce and spinach under 30% reduced irrigation by partially restoring yield, reducing harvest delay, and improving multiple indicators of plant physiological status. Consortium performance varied across crops and treatments, and most treatments did not fully restore yield to the level of fully irrigated plants, but microbial inoculation consistently reduced the severity of water-deficit effects relative to untreated controls. These findings support continued development of multi-species microbial inoculants as scalable tools to enhance crop resilience under increasing water limitation.

## Acknowledgments

The authors thank Dr. Martin Voss, Chief Innovation Officer at Oath Biome, for his advice on the execution of the trial. We are grateful to the microbiology laboratory staff at Oath Biome for producing the microbial inoculants.

## Data availability

The data underlying this article are available via the Zenodo repository DOI:10.5281/zenodo.21111182, https://zenodo.org/uploads/21111182?token=eyJhbGciOiJIUzUxMiJ9.eyJpZCI6IjMxNm QyOGJjLTAzZGYtNDU0Zi05ZjIyLTA2YjUyYjU4OGY4ZSIsImRhdGEiOnt9LCJyYW5kb2 0iOiJjOGZlMzg2MjNkNDQ5ZjY1M2QwZTcyMDI4NTFmOWMwMCJ9.QZqYnl83aOEtlOI FsAEPjTK7HslTsRAl7R-7tABAD4jYDwd1IS-xO7Ekkh6ofwdjTq0crJuaYK8eAxC0Pjc8fg

## Author contribution statement

VB conceived the study. AG, SSB, JLE, JM, AE, JAG compiled, analyzed, and visualized the data for the manuscript. AE, SSB, AG, JM, JAG wrote the initial draft of the manuscript. All authors interpreted the results and contributed to the writing and editing of the manuscript.

## List of Supporting Material Legends

**Figure S1.**
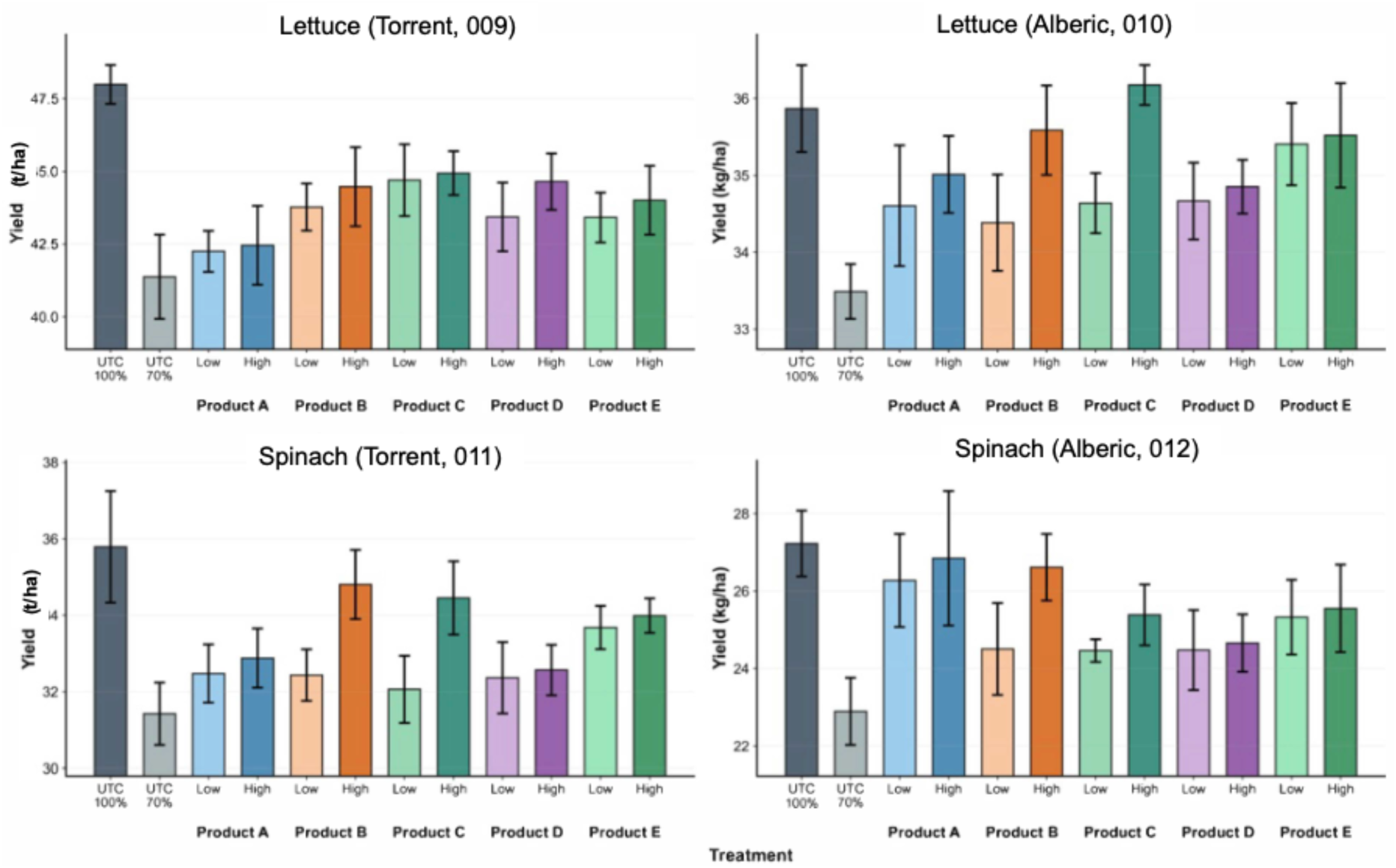
Raw plot-level harvest yield (t ha⁻¹) at each trial site for lettuce and spinach. Bars show treatment means; error bars are standard error of the mean across replicate plots. Data are not spatially detrended. Main-text figures report spatially detrended, across-site analyses with model-based standard errors.

