## Supplemental Table 1 for "Complex microbial consortia improve yield and physiological performance of leafy greens under deficit irrigation"

**Table S1.** Strain-level composition of consortia A–E. Each row is one strain; + indicates inclusion in a consortium. Strain labels distinguish strains within a species.

| Species | Strain | A | B | C | D | E |
|---|---|---|---|---|---|---|
| **Bacteria** | | | | | | |
| *Arthrobacter* sp. AK04 | A | | + | + | + | |
| | B | | | | + | + |
| *Arthrobacter* sp. StoSoilA2 | A | | + | + | + | |
| *Bacillus haynesii* | A | | + | + | + | + |
| *Bacillus inaquosorum* | A | | + | + | + | |
| | B | | + | | | + |
| *Bacillus licheniformis* | A | | + | + | | + |
| *Bacillus paralicheniformis* | A | | | | | + |
| *Bacillus safensis* | A | | + | + | | + |
| | B | | + | + | | |
| *Bacillus spizizenii* | A | | + | | | + |
| *Bacillus subtilis* | A | | + | + | | |
| | B | | + | + | | |
| | C | | | + | | + |
| *Bacillus velezensis* | A | + | | | | |
| | B | | | + | | |
| | C | | | + | | |
| *Kocuria oxytropis* | A | | + | | | + |
| *Lysinibacillus capsici* | A | | + | | + | |
| *Lysinibacillus sphaericus* | A | + | | | | |
| *Microbacterium* sp. MYb45 | A | + | + | + | | |
| *Paenibacillus cellulositrophicus* | A | + | + | + | + | + |
| | B | + | | + | | + |
| *Peribacillus frigoritolerans* | A | | + | + | | + |
| | B | | + | + | | + |
| | C | | | | | + |
| *Priestia aryabhattai* | A | | + | + | + | |
| | B | | | + | | |
| | C | | | | | + |
| *Priestia endophytica* | A | | + | + | | |
| | B | | | | | + |
| *Priestia megaterium* | A | | + | + | | + |
| *Rhodococcus baikonurensis* | A | + | + | + | | |
| *Streptomyces albogriseolus* | A | | + | + | | |
| *Streptomyces bacillaris* | A | + | + | | | |
| | B | + | | | | |
| | C | | + | + | | |
| *Streptomyces bikiniensis* | A | | + | | | |
| | B | | + | | | |
| *Streptomyces eurythermus* | B | | + | | | |
| *Streptomyces fimbriatus* | A | | + | + | | |
| *Streptomyces globosus* | A | + | + | | + | |
| | B | + | + | | | |
| | C | + | | | | |
| | D | + | | | | |

| Species | Strain | A | B | C | D | E |
|---|---|---|---|---|---|---|
| | E | | + | + | | |
| | F | | + | | | |
| | G | | + | + | | |
| | H | | + | + | | |
| | I | | + | + | | |
| | J | | | + | | |
| *Streptomyces griseoaurantiacus* | A | | + | | | |
| | B | | | + | | |
| *Streptomyces griseoincarnatus* | A | + | | | | |
| | B | + | | | | |
| | C | | + | | | |
| *Streptomyces griseoviridis* | A | | + | | | + |
| *Streptomyces misionensis* | A | + | | | | |
| | B | + | + | | | |
| | C | | + | + | | |
| | D | | + | + | | |
| | E | | + | + | | |
| | F | | + | + | + | |
| | G | | + | + | | |
| *Streptomyces murinus* | B | + | + | + | | |
| | C | + | | | | |
| | D | + | | | | |
| *Streptomyces nogalater* | A | + | | | | |
| *Streptomyces olivaceus* | A | | + | | + | |
| *Streptomyces rameus* | A | + | | | | |
| | B | + | | | | |
| | C | | + | | | |
| | D | | + | | + | |
| *Streptomyces rochei* | A | | + | + | | |
| | B | | + | + | | |
| | C | | | + | | |
| | D | | | + | | |
| *Streptomyces sennicomposti* | A | + | + | | | |
| | B | + | | | | |
| | C | + | | | | |
| | D | | + | | | |
| | E | | + | | | |
| *Streptomyces* sp. | A | + | | | | |
| | B | + | | | | + |
| | C | | + | | | |
| | D | + | | | | |
| | E | | + | | | |
| **Arbuscular mycorrhizal fungi (AMF)** | | | | | | |
| *Claroideoglomus etunicatum* | A | | + | + | + | + |
| *Funneliformis mosseae* | A | | + | + | + | + |
| *Glomus aggregatum* | A | | + | + | + | + |
| *Rhizophagus irregularis* | A | | + | + | + | + |
| **Ectomycorrhizal fungi (EMF)** | | | | | | |
| *Pisolithus tinctorius* | A | | + | | | + |

| Species | Strain | A | B | C | D | E |
|---|---|---|---|---|---|---|
| *Rhizopogon amylopogon* | A | | + | | + | + |
| *Rhizopogon fulvigleba* | A | | + | | + | + |
| *Rhizopogon luteolus* | A | | + | | + | + |
| *Rhizopogon villosulus* | A | | + | | + | + |
| *Scleroderma cepa* | A | | + | | | + |
| *Scleroderma citrinum* | A | | + | | | + |
| **Bacterial strains (n)** | | 28 | 55 | 41 | 12 | 19 |
| ***Streptomyces* strains (n)** | | 22 | 34 | 19 | 4 | 2 |
| **AMF species (n)** | | 0 | 4 | 4 | 4 | 4 |
| **EMF species (n)** | | 0 | 7 | 0 | 4 | 7 |

In total, 87 bacterial strains and 11 fungal species inocula were used across consortia A–E.
