## Supplemental Table 2 for "Complex microbial consortia improve yield and physiological performance of leafy greens under deficit irrigation"

**Table S2.** Site, crop and management details of the four greenhouse trials

| Parameter | 009 – Lettuce, Torrent | 010 – Lettuce, Alberic | 011 – Spinach, Torrent | 012 – Spinach, Alberic |
|---|---|---|---|---|
| Trial ID | 009.25-RZM-HYDSTR-LACSA | 010.25-RZM-HYDSTR-LACSA | 011.25-RZM-HYDSTR-SPQOL | 012.25-RZM-HYDSTR-SPQOL |
| Crop | Lettuce (Lactuca sativa) | Lettuce (Lactuca sativa) | Spinach (Spinacia oleracea) | Spinach (Spinacia oleracea) |
| Variety | Romana Cercas | Romana Cercas | Grebe RZ | Grebe RZ |
| Site | Torrent, Valencia | Alberic, Valencia | Torrent, Valencia | Alberic, Valencia |
| Coordinates | 39°23′46″ N, 0°31′11″ W | 39°06′33″ N, 0°32′09″ W | 39°23′46″ N, 0°31′11″ W | 39°07′44″ N, 0°32′30″ W |
| Altitude; slope | 121 m; 0% | 34 m; 1% | 125 m; 0% | 29 m; 2% |
| Trial period | 6 Mar – 30 Apr 2025 | 13 Mar – 25 Apr 2025 | 11 Mar – 30 Apr 2025 | 14 Mar – 24 Apr 2025 |
| ***Soil*** | | | | |
| Sand / silt / clay (%) | 28.5 / 17.3 / 54.2 | 0.7 / 19.9 / 79.4 | 30.3 / 19.6 / 50.1 | 2.2 / 17.4 / 80.4 |
| Texture class | Clay | Clay | Clay | Clay |
| pH | 7.1 | 7.4 | 6.9 | 8.26 |
| Organic matter (%) | 2.9 | 2.9 | 2.9 | 4.5 |
| ***Experimental design and planting*** | | | | |
| Design | RCB, 12 treatments × 6 replicates | RCB, 12 treatments × 6 replicates | RCB, 12 treatments × 6 replicates | RCB, 12 treatments × 6 replicates |
| Plot size | 2 × 5 m (10.0 m²) | 2.5 × 3.996 m (9.99 m²) | 2 × 5 m (10.0 m²) | 2.5 × 3.996 m (9.99 m²) |
| Rows per plot; row spacing | 4; 0.50 m | 4; 0.63 m | 4; 0.50 m | 4; 0.63 m |
| Within-row spacing | 0.33 m | 0.33 m | 0.33 m | 0.33 m |
| Plants per plot; density | 60; 60,606 ha⁻¹ (6.06 m⁻²) | 48; 48,000 ha⁻¹ (4.80 m⁻²) | 60; 60,606 ha⁻¹ (6.06 m⁻²) | 48; 48,000 ha⁻¹ (4.80 m⁻²) |
| Transplanting (by hand) | 5 Mar 2025 | 13 Mar 2025 | 13 Mar 2025 | 14 Mar 2025 |
| ***Greenhouse conditions during the trial*** | | | | |
| Mean daily min – max air T (°C) | 12.4 – 34.9 | 11.4 – 42.4 | 12.4 – 36.1 | 9.7 – 39.7 |
| Absolute min – max air T (°C) | 5.1 – 53.2 | 4.8 – 61.3 | 5.1 – 53.2 | 3.1 – 52.7 |
| Days with max T ≥ 40 °C | 17 of 57 | 32 of 44 | 19 of 49 | 27 of 42 |
| Mean daily min – max RH (%) | 40 – 74 | 39 – 94 | 37 – 73 | 29 – 96 |
| Absolute minimum RH (%) | 16 | 21 | 16 | 12 |
| ***Irrigation and crop management*** | | | | |
| System | Drip, one 2 L h⁻¹ emitter per plant | Drip, one 2 L h⁻¹ emitter per plant | Drip, one 2 L h⁻¹ emitter per plant | Drip, one 2 L h⁻¹ emitter per plant |
| Frequency | 6 d week⁻¹ | 6 d week⁻¹ | 6 d week⁻¹ | 6 d week⁻¹ |
| Establishment (all plots) | 60 min d⁻¹, 6–13 Mar | 60 min d⁻¹, 13–17 Mar | 60 min d⁻¹, 13–18 Mar | 60 min d⁻¹, 14–17 Mar |
| Full irrigation (T1) | 45 min d⁻¹ (1.50 L plant⁻¹) | 40 min d⁻¹ (1.33 L plant⁻¹) | 45 min d⁻¹ (1.50 L plant⁻¹) | 40 min d⁻¹ (1.33 L plant⁻¹) |
| Deficit irrigation (T2–T12) | 30 min d⁻¹ (1.00 L plant⁻¹) | 25 min d⁻¹ (0.83 L plant⁻¹) | 30 min d⁻¹ (1.00 L plant⁻¹) | 25 min d⁻¹ (0.83 L plant⁻¹) |
| Delivered ratio (nominal 70%) | 67% | 62.5% | 67% | 62.5% |
| Deficit onset (DA first appl.) | 14 Mar (8 d) | 18 Mar (5 d) | 19 Mar (5 d) | 18 Mar (4 d) |
| Fertilisation | Base dressing 27 Feb: NPK 15-15-15, 200 kg ha⁻¹ + sheep pellet, 600 kg ha⁻¹; no fertigation | Fertigation 13 Mar, 26 Mar, 9 Apr (see footnote a) | Base dressing 6 Mar: NPK 15-15-15, 200 kg ha⁻¹ + sheep pellet, 600 kg ha⁻¹; no fertigation | Fertigation 13 Mar, 26 Mar, 9 Apr (see footnote a) |
| Weed control (manual tillage) | 13, 20, 27 Mar; 3 Apr | 26 Mar; 7 Apr | 20, 27 Mar; 3, 17 Apr | 26 Mar; 7 Apr |
| ***Inoculant application*** | | | | |
| Application A (BBCH) | 6 Mar 2025 (12) | 13 Mar 2025 (13) | 14 Mar 2025 (13) | 14 Mar 2025 (12) |
| Application B (BBCH) | 20 Mar 2025 (17) | 27 Mar 2025 (16) | 28 Mar 2025 (18) | 28 Mar 2025 (16) |
| Application C (BBCH) | 7 Apr 2025 (33) | 10 Apr 2025 (19) | 11 Apr 2025 (19) | 11 Apr 2025 (43) |
| Interval A→B→C (d) | 14, 18 | 14, 14 | 14, 14 | 14, 14 |

| | | | | |
|---|---|---|---|---|
| Equipment; pressure | Motorised knapsack (Pulmic), SJ303 drench nozzle; 2 bar | Motorised knapsack (Pulmic), SJ303 drench nozzle; 6 bar | Motorised knapsack (Pulmic), SJ303 drench nozzle; 2 bar | Motorised knapsack (Pulmic), SJ303 drench nozzle; 6 bar |
| Application volume | 2300 L $ha^{-1}$ (38 mL $plant^{-1}$) | 2500 L $ha^{-1}$ (52 mL $plant^{-1}$) | 2300 L $ha^{-1}$ (38 mL $plant^{-1}$) | 2500 L $ha^{-1}$ (52 mL $plant^{-1}$) |
| Suspension conc. (250 / 500 g $ha^{-1}$) | 0.109 / 0.217 g $L^{-1}$ | 0.100 / 0.200 g $L^{-1}$ | 0.109 / 0.217 g $L^{-1}$ | 0.100 / 0.200 g $L^{-1}$ |
| Dose per plant (250 / 500 g $ha^{-1}$) | $4.1 \times 10^4$ / $8.3 \times 10^4$ CFU | $5.2 \times 10^4$ / $1.0 \times 10^5$ CFU | $4.1 \times 10^4$ / $8.3 \times 10^4$ CFU | $5.2 \times 10^4$ / $1.0 \times 10^5$ CFU |
| Dose per hectare (250 / 500 g $ha^{-1}$) | $2.5 \times 10^9$ / $5.0 \times 10^9$ CFU | $2.5 \times 10^9$ / $5.0 \times 10^9$ CFU | $2.5 \times 10^9$ / $5.0 \times 10^9$ CFU | $2.5 \times 10^9$ / $5.0 \times 10^9$ CFU |
| ***Harvest*** | | | | |
| Harvest date | 30 Apr 2025 | 25 Apr 2025 | 30 Apr 2025 | 24 Apr 2025 |
| Days after application C | 23 | 15 | 19 | 13 |
| Harvest area (two central rows) | 1.00 × 2.00 m = 2.00 $m^2$ (12 plants) | 1.26 × 1.98 m = 2.49 $m^2$ (12 plants) | 1.00 × 1.98 m = 1.98 $m^2$ (12 plants) | 1.26 × 1.98 m = 2.49 $m^2$ (12 plants) |
| Estimated harvest delay, stressed control (d) | 4 | 6 | 3 | 6 |

a Fertigation at Alberic (trials 010 and 012) delivered ammonium nitrate 33.5-0-0 at 14.2, 185.2 and 85.5 kg $ha^{-1}$, phosphoric acid 0-52-0 at 4.75, 76 and 14.25 kg $ha^{-1}$, and potassium nitrate 13.7-0-46.3 at 21.75, 282.75 and 130.5 kg $ha^{-1}$ on 13 March, 26 March and 9 April 2025 respectively. Rates are those specified per hectare in the fertigation programme; delivery relative to irrigation treatment was not recorded (see Methods).

b Greenhouse air temperature and relative humidity were logged by the contract research organisation as daily minima and maxima only; daily means were not recorded. Values given are the means of the daily minima and of the daily maxima over the period from transplanting to harvest.

c Deficit treatments are designated '70% irrigation' throughout; the ratios delivered relative to the fully irrigated control were 67% at Torrent and 62.5% at Alberic.

d The third application in trial 009 was made 18 days after the second rather than the scheduled 14 days.

e Doses are expressed as colony-forming units, calculated from the nominal formulation titre of $1 \times 10^7$ CFU $g^{-1}$ (lot 04020250101_OSL35, used in all four trials).

f All trials were conducted by GMW Bioscience (Alberic, Valencia, Spain) under Good Experimental Practice (GEP accreditation EOR67/08), following EPPO PP 1/135(4), PP 1/152(4) and PP 1/181(5), and CEN 17700-1 and CEN 17700-3.
